# Optimisation of skeletal muscle sampling for cultured fat production

**DOI:** 10.1101/2025.01.22.634340

**Authors:** Richard G. J. Dohmen, Rui Hueber, Beatriz Martins, Maria Ana Gouveia, Simon Zschieschang, Marjolein de Vries, John Caubergh, Ralph Eussen, Mark J. Post, Joshua E. Flack

## Abstract

To produce cultured meat at a commercially-relevant scale, bioprocesses must be robust, standardised and cost-efficient. In this respect, the limited *in vitro* lifespan of primary cells poses a major challenge, which requires regular cell sourcing by means of biopsy. We have previously shown that muscle-derived fibro-adipogenic progenitor cells (FAPs) represent a promising starting cell type for cultured fat production. These cells proliferate for a high number of population doublings (PDs) in serum-free growth medium. However, with accumulating PDs, FAPs lose their adipogenic differentiation capacity, thereby limiting the amount of cultured fat that can be produced from a given starting sample. Donor animal characteristics, including age, may affect this loss of differentiation. Here, we performed a longitudinal biopsy study to ask whether physiological differences between donor animals are reflected in FAP cell biology, and if this has an effect on loss of FAP differentiation capacity. We sampled twelve Limousin cattle over the first two years of their lives, successfully performing 144 muscle biopsies. Animals younger than six months demonstrated higher FAP yields per gram of muscle tissue, making them ideal donors for cultured fat production, but we observed little difference in the proliferation or differentiation of derived cell cultures. This work also provides valuable insights into the tissue harvesting process, highlighting the need for new, robust and standardised biopsy systems to mitigate contamination risk.

## Introduction

Cultured meat (also known as ‘cultivated meat’) biotechnologies offer the potential to alleviate the environmental and animal welfare issues associated with conventional animal agriculture^1,2^. Cells are harvested from biopsied tissue samples and cultured *in vitro* to produce edible skeletal muscle tissue for human consumption^1,3,4^. However, to reach the large-scale production required for commercialisation, numerous challenges remain to be overcome, including the identification of optimal stem cell donors and the standardisation and automation of cell banking^1,5,6^.

Alongside multinucleated myofibers, skeletal muscle comprises inter- and intramuscular fat (IMF) tissue (also referred to as ‘marbling’), a key contributor to its palatability^7,8^. IMF is derived from tissue-resident interstitial cells called fibro-adipogenic progenitors (FAPs)^9–12^, which can be isolated from tissue samples concurrently with satellite cells (SCs)^13^. FAPs proliferate robustly, achieving a large number of population doublings (PDs) with a near-constant growth rate^13,14^. Moreover, their mature differentiation results in cultured fat tissue that visually resembles traditional fat and closely mimics it with respect to both triglyceride composition^13,14^ and taste^13^. However, FAPs invariably lose their adipogenic differentiation capacity over the course of proliferation^13^. Characteristics of the donor animal may affect the rate of this loss of differentiation^15,16^, highlighting the importance of identifying optimal donor animals^4,5^. Numerous donor characteristics can affect IMF deposition^17–19^. For instance, differences in IMF and collagen deposition have been found in anatomically distinct muscles^20,21^, highlighting the importance of the site of biopsy. Other factors that influence IMF deposition are sex^19,22–24^, breed^18,25–27^ and age^28,29^. Differences in bovine marbling are partially the result of FAP hyperplasia^30–32^, which mainly occurs up to 250 days after birth (see review^33^). FAP numbers per tissue volume decline due to hypertrophy^33^ and as observed in mice, as animals age^34,35^. However, whether these physiological differences correspond to variations in FAP cell biology, and how that might affect the loss of differentiation capacity *in vitro*, is largely unknown.

Here, we conducted a longitudinal biopsy study, in which we sampled the same animals at different stages of development. This approach allowed us to simultaneously address the effect of donor age, sex and site of biopsy, on FAP yield, proliferation rate and adipogenic differentiation capacity. We successfully performed 144 muscle biopsies from twelve Limousin cattle (six male, six female), at six timepoints in the first two years of their lives, from two distinct skeletal muscles (semitendinosus and brachiocephalicus). We observed frequent microbial contamination related to biopsy sampling, reflecting the challenges of maintaining aseptic conditions during sampling in real life farm conditions, and highlighting potential improvements for biopsy systems. This work provides insight into the sourcing of primary cells for immediate use in cultivated meat applications, or as the starting point for development of optimised cell lines.

## Results

### Study design, donor animal characteristics and incisional biopsy process

To investigate the effect of donor characteristics on the yield, proliferation and adipogenic differentiation of FAPs, we performed skeletal muscle biopsies of the semitendinosus and brachiocephalicus of twelve Limousin cattle at six timepoints over the course of two years (Figs. 1a, b). Animals rapidly gained weight over the timecourse (although between 12 to 18 months there was no statistical significant increase), irrespective of sex. Sampling was performed using incisional biopsies, which whilst more invasive than suction-assisted needle biopsies^4^, was far superior in terms of ease and yield of sampling. Although we successfully obtained 144 incisional biopsies, we observed a large variability in sample mass between biopsies, though without a significant difference between timepoint, sex or muscle type (Fig. 1b).

**Figure 1:**
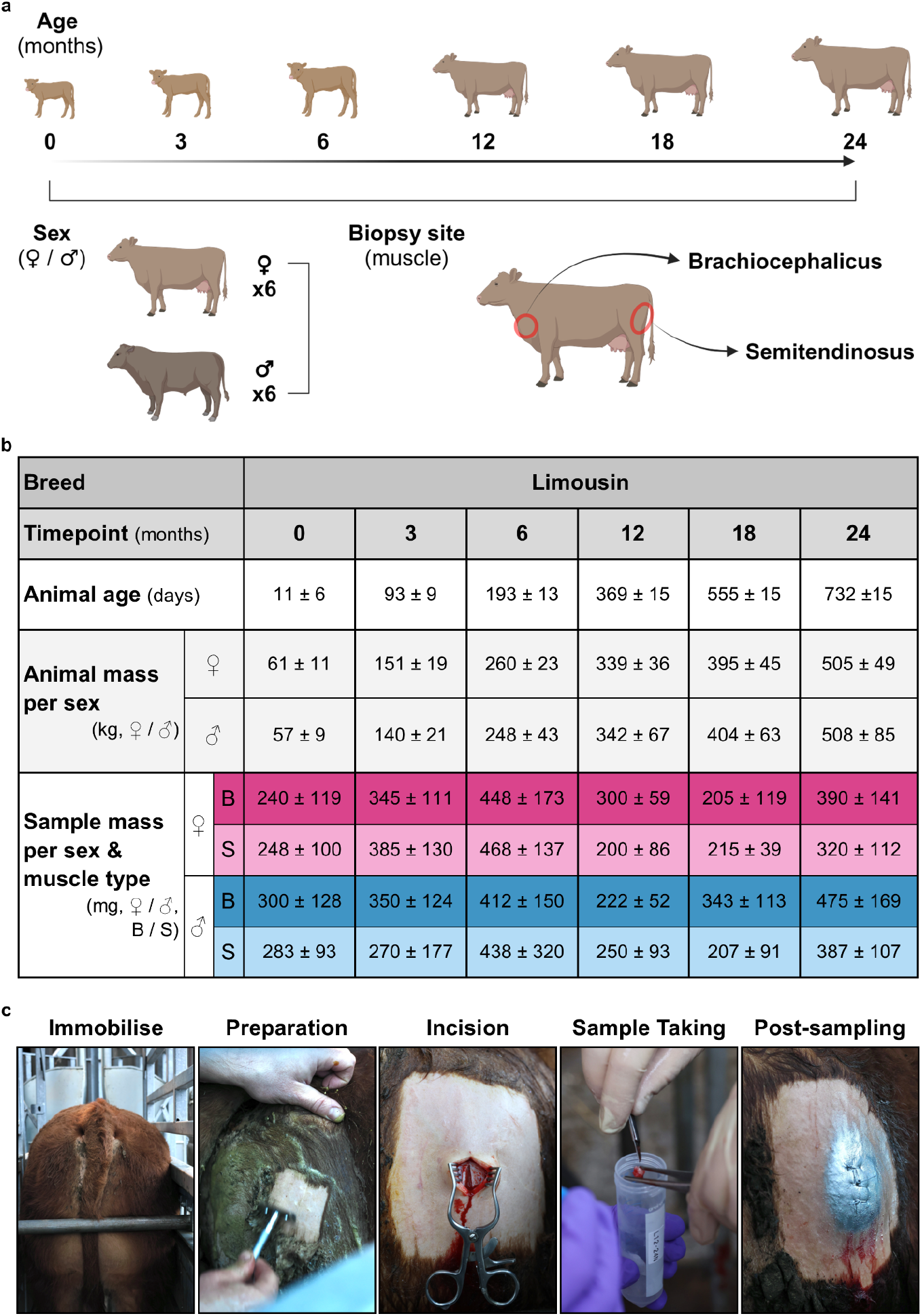
Overview of study design, animal characteristics and incisional biopsy process. **a** Overview of experimental design of the longitudinal study. **b** Key phenotypic characteristics of Limousin age, mass and sample per timepoint at the moment of sampling, compared in this study. Sex is represented as follows: Female ♀ and Male ♂, and muscle type: Brachiocephalicus (B) and Semitendinosus (S). All data is shown as mean ± sd (n = 12 for animal age, n = 6 for animal and sample mass). **c** Representative photographic images of key steps within the biopsy process.

Once enzymatically digested, we employed a pre-culture phase to enrich our cell cultures for the two main muscle-derived stem cell types of interest (FAPs and SCs). During this phase, we observed microbial (bacterial or fungal) contamination in 30% (42/144) of cultures. The majority of contaminations were observed within two to four days (with two incidents during subsequent stages of culture) corresponding to the point where antibiotic concentration was reduced. Despite increased washing of the biopsied tissue and stringent aseptic handling, contaminations still occurred, suggesting that they arise due to exposure to the non-sterile environment after incision (Fig. 1c), and pointing to the need for a fully closed biopsy system (see Discussion).

### FAP yield decreases with donor age

Next, we aimed to determine the yield of FAPs isolated per volume of tissue. Quantification of cell numbers immediately post-isolation is complicated by the presence of structural and cellular debris related to enzymatic digestion, so we chose to quantify cells at the end of the pre-culture phase. We observed empirically that yields were maximised when cultures approached confluence in the pre-culture phase. This meant that the length of this phase differed between biopsies (ranging from two to thirteen days), with the majority of samples having five to seven days of pre-culture (Fig. 2a, left panel). Sample mass and FAP yield at the end of pre-culture showed similar distribution (Fig. 2a, middle and right panels, respectively). Both sample mass and FAP yields at the end of pre-culture varied significantly, reflecting the inherent variability in biopsy sizes and tissue quality. The majority of samples weighed between 200-499 mg (mean ± standard deviation (sd): 335 ± 152 mg) and yields ranged from 80,000 to more than 3M (mean ± sd: 2.74 ± 2.6M). Sample mass negatively correlated with length of pre-culture (R = -0.2215, *P* = 0.0253), and positively correlated with FAP yields (R = 0.4567, *P* = <0.0001), suggesting that measured yields at the end of pre-culture reasonably reflect actual counts per given biopsy volume (Fig. 2b, left and right panel, respectively).

**Figure 2:**
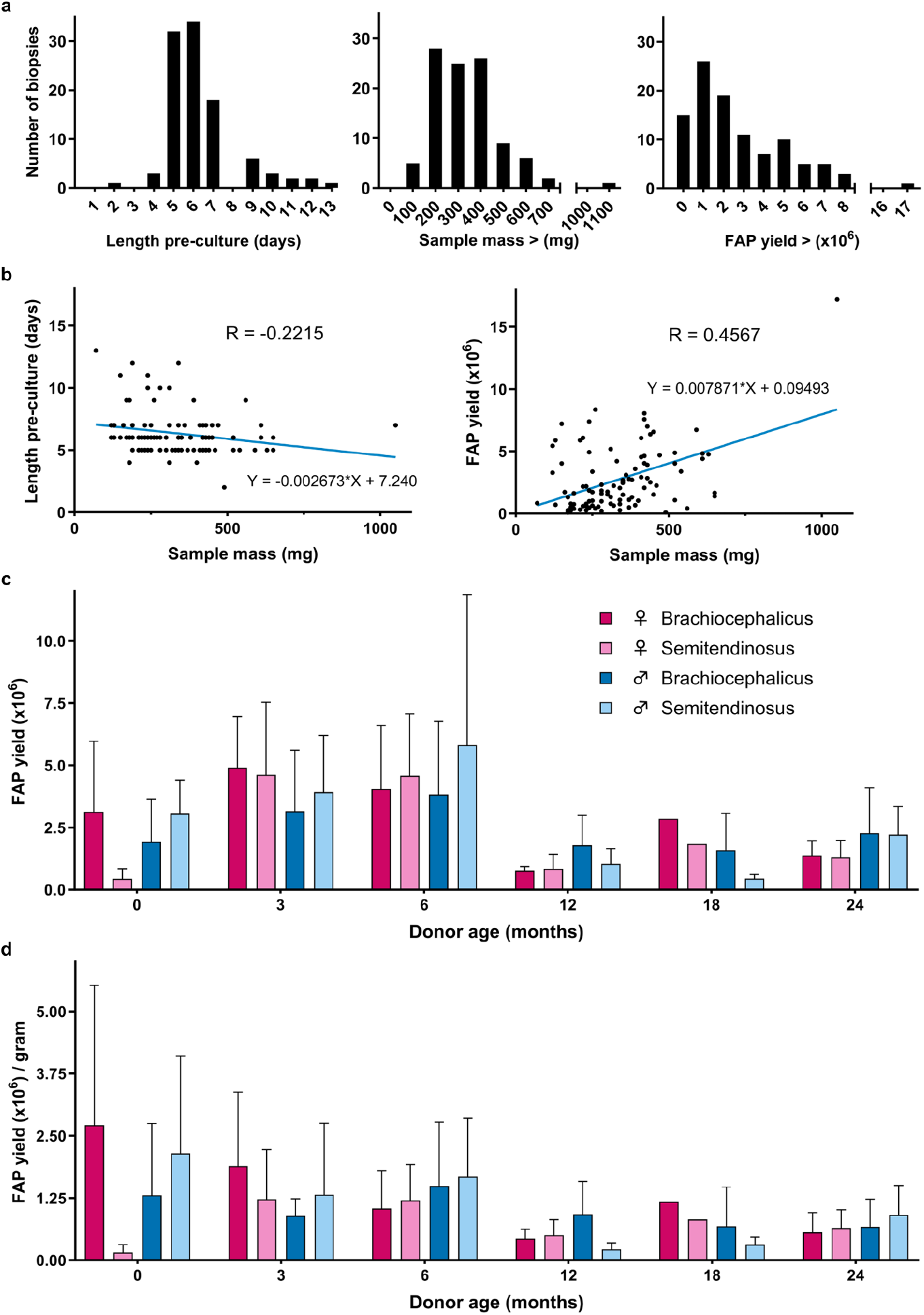
FAP yield decreases with donor age. **a** Histograms showing the distribution frequency of 102 samples in length of pre-culture, sample mass and FAP yield at end of pre-culture. Bins represent days, or ranges of mass or yield (left, middle and right panel, respectively). **b** Scatter plots of 102 samples showing Pearson correlation of sample mass with length of pre-culture or FAP yield (left or right panel, respectively), trendline with equation and correlation coefficients are indicated. **c** FAP yields at end of pre-culture, and **d** FAP yields per gram per condition over the six timepoints (0, 3, 6, 12, 18 and 24 months). Bars represent conditions, where colour represent sex (pink = Female (♀), blue = Male (♂)) and shade represents muscle type (darker = Brachiocephalicus, lighter = Semitendinosus). Data is shown as mean ± sd (n = 1-6 per condition and timepoint).

Comparing FAP yields at the end of the pre-culture phase across donor ages, sex and muscle type revealed a distinct drop in FAP yields between six and twelve months of age. Before and after this drop, yields were reasonably constant (Fig. 2c), in line with prior observations on muscle hyperplasia and hypertrophy in cattle^33^. Age was the only significant explanatory variable of FAP yield (F (2.240, 22.40) = 7.310, *P* = 0.0028). We estimated the FAP yields per gram of tissue, using the yield at the end of pre-culture, the duration of pre-culture and assuming FAPs proliferated at 0.5 PDs per day (a previously observed average). We determined FAP yields per gram by dividing the calculated FAP yields at the start of pre-culture by sample mass, thus defining FAP yield per gram. After normalisation, however, no significant effect of any of the independent variables or their interactions on FAP yield was observed (Fig. 2d), most likely variability was influenced by assumptions in the PDs per day during pre-culture. Irrespective, these data suggest that animals younger than (at least) 6 months are ideal donors in terms of FAP yield. Within these 6 months, the same animal could also be sampled at least three times without differing FAP yields, increasing the amount of cultured fat that could be produced from a given donor.

### Donor age has no effect on FAP proliferation and differentiation capacity

To assess the effect of donor age on FAP proliferation and differentiation capacity, we performed a long-term proliferation experiment in serum-free growth medium (SFGM) with FAPs derived from semitendinosus biopsies at four distinct timepoints (0, 3, 6 and 12 months) from four donor animals. Over the course of 52 days, cultures proliferated for 25 cumulative population doublings (PDs), with no significant difference between donor ages (Fig. 3a, left). Cultures proliferated at a similar rate, with an average of 0.51 ± 0.12 PDs per day and no significant difference between ages (Fig. 3a, right). During the course of these cultures we did not observe differences in cell morphology between donor ages, but with increased cellular age, FAPs flattened and enlarged (Supplementary Fig. 1).

**Figure 3:**
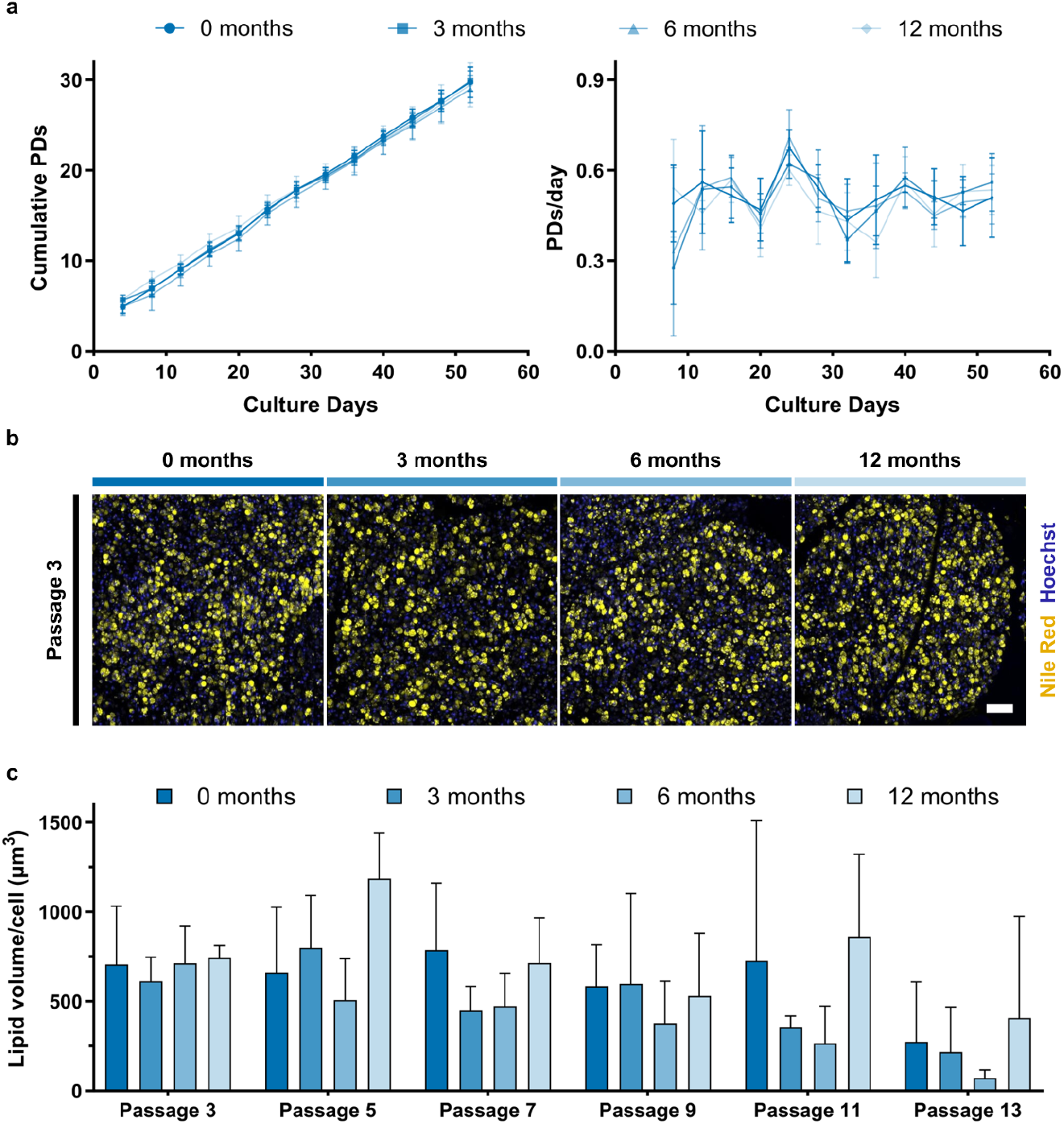
Donor age has no effect on FAP proliferation and differentiation. **a** Growth curve (left panel, showing cumulative population doublings) and growth rate (right) of FAPs grown in (SFGM) for thirteen passages. Different shaded lines and symbols represent donor age, sd is indicated (n = 4). **b** Representative maximum intensity projection confocal microscopy images of 28 day differentiated beads corresponding to samples quantified in **c**. Yellow = Nile Red, blue = Hoechst, scale bar = 100 μm. **c** Lipid volume per cell of 28 days differentiated beads generated from FAPs that had undergone proliferation in SFGM conditions in **a**. Data is shown as mean ± sd (n = 4).

We also assessed the effect of donor and cellular ageing on adipogenic differentiation. FAPs were encapsulated in alginate hydrogel and extruded in a drop-wise fashion to produce adipogenic ‘bead’ constructs. We previously observed that FAP differentiation dropped significantly with accumulating PDs^13^. In this experiment, adipogenesis was not reduced significantly with increasing cellular age, though this may be related to the relatively low fat volumes per nucleus observed throughout the experiment (ranging between 100 to 1000 µm^3^). We also observed significant intra- and inter-donor variability in fat volume per nucleus within and between timepoints, further complicating the data interpretation and highlighting the need for a more robust differentiation assay. More importantly, however, there was no significant difference in adipogenic differentiation in cells derived from donors of different ages. This suggests that the reduction of adipogenic differentiation due to *in vitro* ageing occurs at a similar rate for FAPs derived from 0, 3, 6 and 12 month old donors.

## Discussion

An efficient bioprocess for production of cultured meat from biopsies from live animals requires the optimisation of many parameters, including stem cell yield and the ability of the derived cells to robustly proliferate and differentiate at higher population doublings. While many early cultured meat products are formed from undifferentiated cell-based ingredients, FAPs represent a promising cell type for the production of genuine, differentiated cultured fat^13^. However, it is unclear whether and how physiological characteristics of stem cell donor animals influence FAP proliferation and differentiation *in vitro*. Here, we launched a longitudinal biopsy study to investigate the effects of age, sex and biopsy site on FAP yield, proliferation and differentiation.

Although tissue sampling is common practice in human and veterinary medicine, cell-based bioprocesses have different requirements. Maintaining total sterility is critical, as microbial contamination will compromise subsequent cell culture. Achieving such standards is particularly challenging on a farm, where animal and environmental conditions are harder to control than in a clinical setting. Additionally, cell-based processes require obtaining as large a quantity of starting material as possible in a standardised manner through a minimally invasive tissue sampling procedure^5^. Suction-assisted needle biopsies meet some of these requirements, but are not optimised for tough tissues or the harvesting of viable cells, and in our pilot experiments we were unable to obtain meaningful numbers of cells in this fashion. Here, using an incisional biopsy protocol, we were able to successfully obtain 144 muscle samples from twelve animals over a period of two years. Not surprisingly, incisional biopsies resulted in higher average sample mass compared to reported human vacuum- or suction-assisted needle biopsy yields, although the amount of tissue obtained was also variable^36,37^. To minimise muscle injury, sample mass was substantially lower than suggested maximal yields^4^ and only slightly higher than human (automated) vacuum-assisted needle biopsy^36^, losing its advantage over needle biopsies. Considering its invasiveness and the high risk of contamination due to air exposure, incisional biopsies may not be the optimal method for initiating a bioprocess.

Future efforts should thus focus on developing needles to perform aseptic and standardised tissue sampling for a cultured meat bioprocess. Most biopsy needles are designed for human medicine, favouring smaller bore sizes to minimise trauma. Increasing the size of the needle would allow for more tissue to be collected. A needle comprising an outer needle diameter of 9 mm, combined with an inner needle sample notch length of less than 2 cm, would suffice to obtain ∼0.5 g of muscle tissue, yielding a larger sample mass than traditional diagnostic needles^36,37^. For tissue sampling, spring-loaded systems offer an easy-to-use and consistent alternative over suction- and vacuum-assisted needles^38^. Needle tips should also be compatible with ultrasound guidance to avoid major blood vessels. Lastly, tissue samples should be retrievable inside a canister filled with medium for optimal cellular survival. This novel design would also facilitate the implementation of a fully-enclosed system, minimising contamination risks. Such a proposed system solves aforementioned problems, whilst enhancing efficiency and reproducibility of biopsy sampling in the context of cultured meat production (Fig. 4).

**Figure 4:**
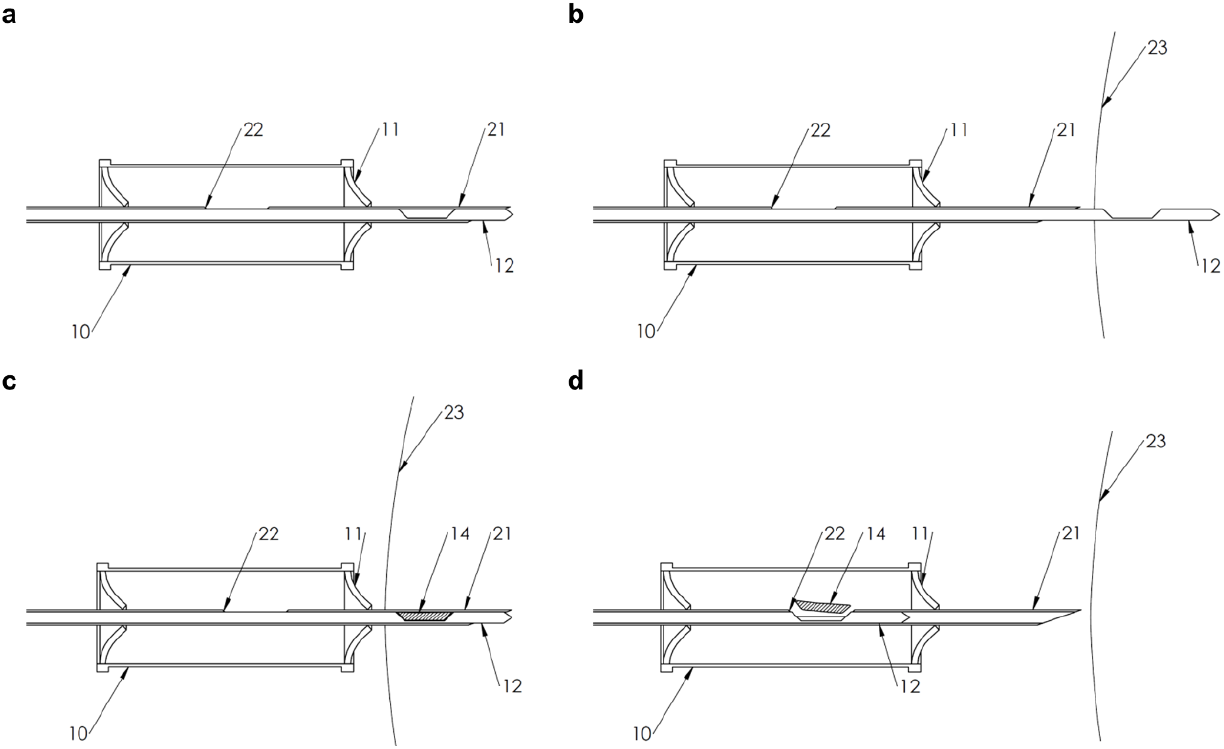
Schematics of a fully-enclosed biopsy system. **a** Biopsy system containing an inline collection vessel (10) primed for sampling. **b** Inner needle (12) shoots forward into the animal (23). **c** Followed by the outer cannula (21) that cuts the sample (14). **d** The inner needle is retracted and releases the sample (14) into an inline collection vessel (10) via an opening (22) in the inner needle (12). This vessel is sterile and contains cell culture medium.

Donor age might affect cultured fat bioprocesses via an effect on FAP yield. Despite the weight gain of all animals, FAP yields after pre-culture only decreased from six to twelve months (Fig. 2). This signifies that even though animals grow and muscles undergo hypertrophy, FAPs proliferate in the first six months. The subsequent drop in FAP yield aligns with the reported end of FAP hyperplasia, which occurs at eight months after birth (see review^33^). This suggests that in terms of FAP yield, animals of up to six months make for good stem cell donors, as starting with larger number of cells requires fewer PDs to create commercially relevant quantities of meat. It would also be interesting to establish FAP yield and behaviour over the full lifetime of an animal. This would avoid sampling animals at an age at which FAP yields and their proliferation and differentiation capacity are unsuitable to start a cultured fat bioprocess. Sectioning of muscle biopsies and immunohistochemistry for known FAP markers might ensure more reliable estimates of FAP numbers per tissue volume, and would also inform potential cell losses during isolation.

Although FAP yield is important, achieving longer proliferation phases (to higher PDs) whilst maintaining robust adipogenic differentiation potential is much more critical for cultured fat bioprocess design, due to the exponential nature of the proliferation process. We observed no differences in FAP proliferation rate or adipogenic differentiation between donors of different ages (Fig. 3). However, we only compared donor ages within the first twelve months of birth. It would be interesting to include older animals to establish an age at which FAP *in vitro* proliferation and adipogenic differentiation is affected, as has been reported for aged murine FAPs^34^. Aside from donor age, it is possible that we failed to capture differences in proliferation and differentiation because of our study design, and future work should extend the proliferation timecourses past 30 PDs. We have previously observed that FAPs from certain donor animals lose their proliferative capacity at higher PDs, ranging from 40 to 50 (unpublished data), which may be an effect of donor age. Our analysis also revealed no significant differences in FAP yield between donor sexes, nor between the two skeletal muscles we sampled (Fig. 2). This may be partially due to our limited sample size and inherent variability in the current sampling technique, but overall our results suggest that donor physiology is not fully recapitulated in the cell biology of the derived cultures. Future research could focus on FAP yield between anatomically distinct muscles with more substantial differences in IMF content.

Finally, improvements in the extent and reproducibility of our adipogenic differentiation constructs, including further optimisation of differentiation medium and the extrusion process for the cell-alginate mixture, could help to reduce intra-donor variability in observed differentiation, and help to reveal more subtle differences between cells from different donors. Similarly, suboptimal proliferation conditions might contribute to the loss of adipogenic differentiation potential *in vitro*. One interesting follow-up would be to culture cells on substrates with lower stiffnesses (e.g. 5 kPa), which might help to extend adipogenic differentiation potential^39^, and allow a more sensitive comparison of cells from different donors. An alternative approach could be to use cell engineering techniques to optimise phenotypic characteristics of FAPs (or other adult stem cell types) such as proliferative lifespan^40,41^.

In conclusion, this work demonstrates that biopsies of cattle aged up to twelve months represent a good starting point for the isolation of FAPs for production of cultured fat, given the lack of difference in proliferation and differentiation between cells from different donor ages. Key challenges remain, including ensuring sterility and standardisation related to biopsy sampling, and we propose that a fully-enclosed biopsy system can address such issues. Additionally, it remains critical for the feasibility of cultured meat bioprocesses to maintain robust differentiation potential after long proliferation phases, something that may be achieved through further process optimisation, or cell line engineering efforts.

## Material & methods

### Experimental design and sample collection

The experiment was performed under an animal experiment licence approved by local and national authorities (AVD1070020209364). All muscle biopsies were taken by a licensed veterinarian. Biopsies of semitendinosus and brachiocephalicus muscles were obtained from twelve Limousin cattle (six males, six females), on each of the six timepoints (0, 3, 6, 12, 18 and 24 months). Housing conditions were similar for all animals, and kept in local pastures (Mar-Oct) or barn (Nov-Feb) with *ad libitum* access to feed and water. All donor animals were part of the conventional meat production system.

For the biopsies, animals were immobilised in an animal handling cage (Cattle crusher A8000, Patura) and sedated with Xylazine. The biopsy site was then washed, shaved, and local anaesthesia (Procamidor) was applied. For sampling, the biopsy site was disinfected (70% ethanol), and a small incision was made piercing skin and fascia. A wound spreader was introduced, and incisional biopsies obtained using tweezers and scalpel. Post sampling, skin was sutured (PGA 6/0), covered with aluminium spray and animal received analgesic (Novem 20) and antidote (Alzane) (Fig. 1c). Post-biopsy check ups were performed daily for ten days. Biopsy samples were transported to the laboratory on ice, in accordance with Dutch national guidelines on handling of animal material.

### Muscle-derived stem cell isolation

SCs and FAPs were isolated from muscle biopsies as previously described, with minor adjustments^13^. Briefly, biopsies were dissociated with collagenase (CLSAFA, Worthington; 45-60 min, 37 °C), filtered through a 40 μm cell strainer, and incubated in ammonium-chloride-potassium (ACK) erythrocyte lysis buffer (1 min, room temperature (RT)). Muscle-derived cells were resuspended in ‘pre-sorting’ serum-free growth medium (ps-SFGM, Supplementary Table 1), and plated on bovine fibronectin (4 μg/cm^2^; Sigma-Aldrich, F1141) coated flasks.

### Fluorescence-activated cell sorting (FACS)

Upon reaching confluence, percentage of two main muscle-derived cell populations (FAPs and SCs) was determined using antibodies against ITGA5 (PE) and ITGA7 (APC) on MACSQuant10 Flow Analyzer (Miltenyi). Based on the percentage of SCs present in the total cell fraction, cells were either sorted or cultured in FAP SFGM (Supplementary Table 1), in case the percentage of SCs was <10%. Cells were sorted on a MACSQuant Tyto Cell Sorter (Miltenyi) using antibodies as described above. Cells were gated into two main populations ITGA5+/ITGA7- (FAPs) and ITGA5-/ITGA7+ (SCs), and sorted into the positive (SCs) and negative (FAPs) collection chambers.

### FAP cell culture

FAPs were cultured on collagen-coated (1.2 μg/cm^2^; bovine skin collagen; C2124, Sigma-Aldrich) flasks in SFGM (Supplementary Table 1) at 5 × 10^3^ cells/cm^2^, and passaged upon reaching confluency (3 to 4 days). Brightfield microscopy images were captured with an EVOS M5000 microscope (Thermo Fisher).

### Adipogenic differentiation bead cell culture

FAPs were resuspended in 0.5% high viscosity non-functionalized alginate solution (Sigma-Aldrich, W201502) at 3 × 10^7^ FAPs/mL. Suspension was extruded into gelation buffer (66 mM CaCl_2_, 10 mM HEPES) in a droplet fashion. Beads were washed, transferred to 48-well tissue culture plates containing adipogenic differentiation medium (Supplementary Table 2)^42^ and incubated on a shaking platform (90 rpm, 37 °C, 5% CO_2_) for 28 days. Medium exchanges were performed every 7 days.

### Immunofluorescence for adipogenic beads

Beads were fixed (4% formaldehyde in gelation buffer; 1 h, RT), and stained overnight at 4 °C (1:500 Nile Red (N1142, Thermo Fisher), 1:625 Hoechst 34580 (63493, Sigma-Aldrich)). Images were captured on a confocal microscope (Stellaris 5, Leica Microsystems) using a 10×/0.4 NA objective lens with 2× zoom and 4 μm Z-steps. Nuclei counts and lipid volumes were quantified using a custom python script for twentyfive tiles per image per sample (n=4 per condition). The total quantified lipid volume was divided by the number of nuclei to calculate lipid volume per cell.

## Statistical analysis

Statistical analysis was performed using Prism 10.0.2 (GraphPad). Analyses involving three or more groups with two or three independent variables (age, sex, biopsy site) were performed using a (mixed-effects) two-way or three-way ANOVA, respectively, with Tukey’s multiple comparison test used for post-hoc comparisons (Fig. 1b, Fig.2c, d, Fig. 3a, c). To account for violations of sphericity, Geisser-Greenhouse correction was applied (Fig.2c, d. Fig. 3c). Pearson correlations of indicated independent and dependent variables were computed assuming a normal distribution (Fig. 2b).

## Data availability

Further data supporting the findings of this study are available from the authors on request.

## Code availability

Scripts for image analysis are available from the authors on request.

## Acknowledgements

We would like to thank the staff of Maatschap Caubergh-Caubergh for supporting this project. We would also like to thank the scientific staff of Mosa Meat B.V. for helpful discussions throughout the course of the experimental work and manuscript writing.

## Author Contributions

RGJD, RH, BM, MAG, SZ and MdV performed experiments and analysis. JC and RE performed animal husbandry and harvested the skeletal muscle biopsies. MJP and JEF supervised the study. RGJD and JEF wrote the manuscript, with input from all authors.

