## Supplementary Material for "Optimisation of skeletal muscle sampling for cultured fat production"

### Supplementary nformation

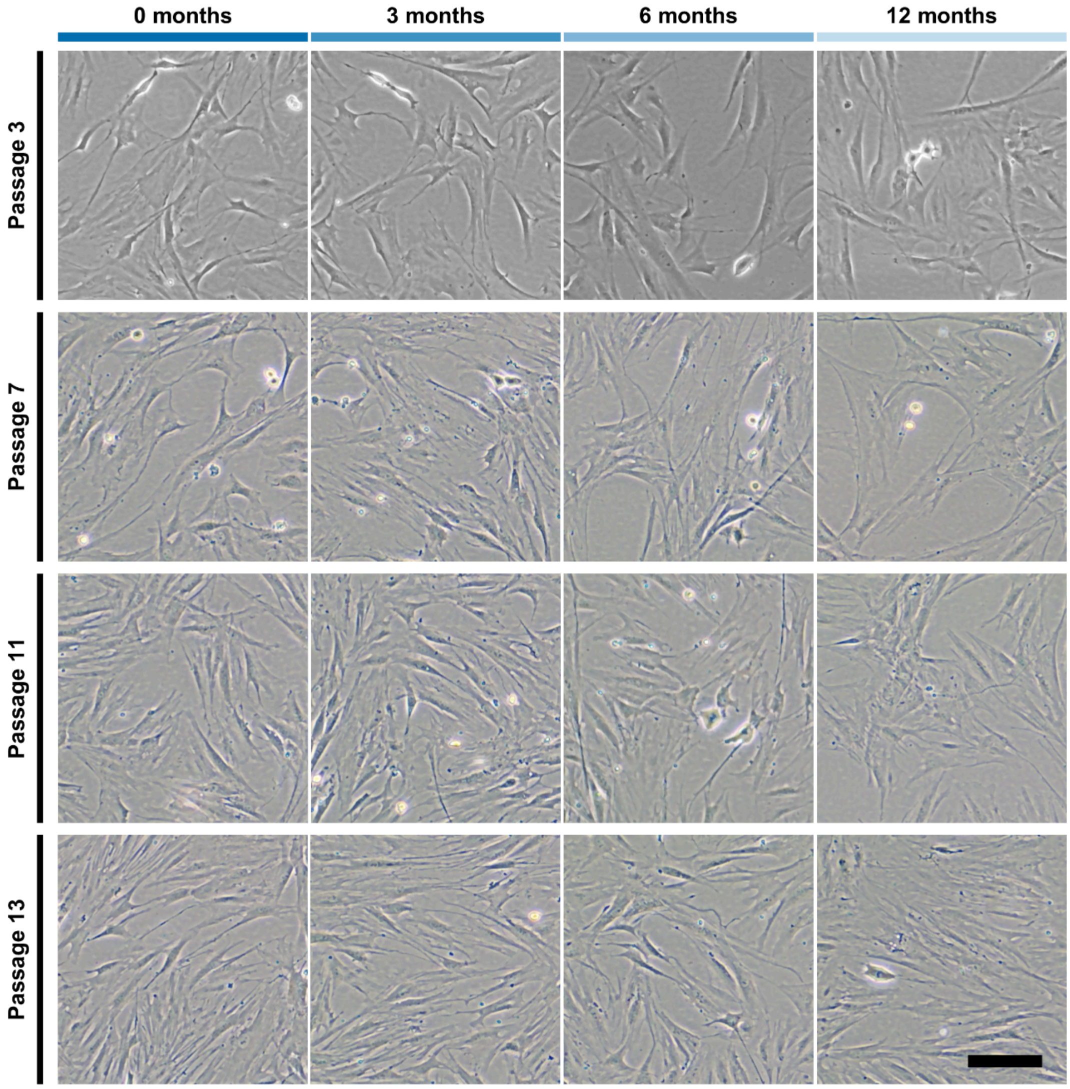


**Supplementary Figure 1: Microscopy images of FAPs at different donor and *in vitro* ages.**

Brightfield microscopy images of sorted FAPs of the same donor at different donor ages (0, 3, 6 and 12 months) and *in vitro* ages (passages 3, 7, 11 and 13). Scale bar = 100 µm.

**Supplementary Table 1:** Serum-free growth media formulations

| # | Component | Reference | Concentration |
|---|---|---|---|
| **Pre-sorting serum-free growth medium (ps-SFGM)** | | | |
| 1 | DMEM/F-12 | 21331-020, Gibco | |
| 2 | α-linolenic acid | L2376, Sigma Aldrich | 1 µg/ml |
| 3 | FGF-2 | 100-18B, Peprotech | 10 ng/ml |
| 4 | Human Serum Albumin | Rc HA NW20, Laurus Bio | 5 mg/ml |
| 5 | HGF | 100-39H, Peprotech | 5 ng/ml |
| 6 | Hydrocortisone | H0135, Sigma Aldrich | 36 ng/ml |
| 7 | IGF-1 | 100-11, Peprotech | 100 ng/ml |
| 8 | IL-6 | 200-06, Peprotech | 20 ng/ml |
| 9 | ITSE | 00-101, biogems | 1% |
| 10 | GlutaMax | 35050-061, Gibco | 1% |
| 11 | Glucose | G7021, Sigma Aldrich | 17.7 mM |
| 12 | L-ascorbic acid 2-phosphate (Vitamin C) | A8960, Sigma Aldrich | 155 µM |
| 13 | PDGF-BB | 100-14B, Peprotech | 10 ng/ml |
| 14 | Penicillin/Streptomycin/ Amphotericin (PSA) | 17-745E, Lonza | 3% |
| 15 | VEGF | 100-20 Peprotech | 10 ng/ml |
| **FAP serum-free growth medium (SFGM)** | | | |
| 1 | DMEM/F-12 | 21331-020, Gibco | |
| 2 | α-linolenic acid | L2376, Sigma Aldrich | 1 µg/ml |
| 3 | FGF-2 | 100-18B, Peprotech | 10 ng/ml |
| 4 | Human Serum Albumin | Rc HA NW20, Laurus Bio | 5 mg/ml |
| 5 | Hydrocortisone | H0135, Sigma Aldrich | 36 ng/ml |
| 6 | IGF-1 | 100-11, Peprotech | 100 ng/ml |
| 7 | IL-6 | 200-06, Peprotech | 20 ng/ml |
| 8 | ITSE | 00-101, biogems | 1% |
| 9 | GlutaMax | 35050-061, Gibco | 1% |
| 10 | Glucose | G7021, Sigma Aldrich | 17.7 mM |
| 11 | L-ascorbic acid 2-phosphate (Vitamin C) | A8960, Sigma Aldrich | 155 µM |
| 12 | PDGF-BB | 100-14B, Peprotech | 10 ng/ml |
| 13 | PSA | 17-745E, Lonza | 1% |

**Supplementary Table 2:** Serum-free adipogenic differentiation medium formulation

| # | Component | Reference | Concentration |
|---|---|---|---|
| **FAP serum-free differentiation medium (DMAD[44])** | | | |
| 1 | DMEM/F-12 | 21331-020, Gibco | - |
| 2 | PSA | 17-745E, Lonza | 1% |
| 3 | HEPES | H3375, Sigma Aldrich | 4.9 mM |
| 4 | Lipid concentrate | 11905031, Thermo Fisher | 0.1% |
| 5 | Hydrocortisone | H0135, Sigma Aldrich | 25 nM |
| 6 | Putrescine | 51799, Sigma Aldrich | 57 µM |
| 7 | Progesterone | P8783, Sigma Aldrich | 17.8 nM |
| 8 | Calcium Chloride | C3881, Sigma Aldrich | 1 mM |
| 9 | L-Ascorbic acid 2-phosphate (Vitamin C) | A8960, Sigma Aldrich | 227 µM |
| 10 | Glucose | G7021, Sigma Aldrich | 17 mM |
| 11 | Insulin | P-2701000, Pan Biotech | 10 µg/ml |
| 12 | BMP4 | 120-05ET, Peprotech | 2 ng/ml |
| 13 | Indomethacin | I7378, Sigma Aldrich | 50 µM |
